# Predicting viral sensitivity to antibodies using genetic sequences and antibody similarities

**DOI:** 10.1101/2025.08.08.669352

**Authors:** Kai S. Shimagaki, Gargi Kher, Rebecca M. Lynch, John P. Barton

## Abstract

For genetically variable pathogens such as human immunodeficiency virus (HIV)-1, individual viral isolates can differ dramatically in their sensitivity to antibodies. The ability to predict which viruses will be sensitive and which will be resistant to a specific antibody could aid in the design of antibody therapies and help illuminate resistance evolution. Due to the enormous number of possible combinations, it is not possible to experimentally measure neutralization values for all pairs of viruses and antibodies. Here, we developed a simple and interpretable method called grouped neutralization learning (GNL) to predict neutralization values by leveraging viral genetic sequences and similarities in neutralization profiles between antibodies. Our method compares favorably to state-of-the-art approaches and is robust to model parameter assumptions. GNL can predict neutralization values for viruses with no observed data, an essential capability for evaluating novel viral strains. We also demonstrate that GNL can successfully transfer knowledge between independent data sets, allowing rapid estimates of viral sensitivity based on prior knowledge.

## Introduction

Viral pathogens such as HIV-1 exhibit extraordinary genetic diversity. Consequently, distinct viral strains can differ dramatically in their sensitivity to antibodies. Predicting whether a given virus is sensitive or resistant to a particular antibody has therefore become a key goal with multiple applications. Antibodies have been used as therapeutic treatments ^1–3^ and for passive immunization ^4^, which can fail when viruses develop resistance ^5–7^. Thus, predicting viral sensitivity or resistance before treatment could enable the rational design of more effective antibody treatment regimens ^8^. Such predictions would also reveal trends in antibody resistance over time, which could inform vaccine design for HIV-1 or other rapidly evolving pathogens.

Experiments alone are insufficient to fully characterize the virus-antibody sensitivity landscape. The space of functional virus sequences is so large that it is effectively unlimited, precluding exhaustive tests even for a single antibody. In response, a variety of computational methods have been developed to attempt to learn patterns of sensitivity and resistance from finitely sampled data ^9^.

For HIV-1, traditional machine learning approaches have used neutralization data coupled with HIV-1 envelope amino acid sequences to predict virus-antibody neutralization values. Support vector machines ^10^ and gradient boosting machines ^11^ have both been used to make binary predictions of sensitivity or resistance. Subsequent work combined additional features (e.g., geographic information for viral isolates) into a “super learning” framework to produce more precise predictions of neutralization values ^12–14^. Methods have also incorporated virus-antibody structural data into a multi-layer perceptron model for resistance prediction, which predicts both binary sensitive/resistant states and IC50 values ^15^.

In recent years, deep learning methods have also been applied to this problem. One approach used recurrent neural networks to capture long-range dependencies in viral sequences ^16^, which was later extended to incorporate attention mechanisms that identify the most relevant amino acid positions for neutralization prediction ^17^. Notably, the latter study incorporated antibody sequence data into the predictive model. Language models have also been applied for resistance prediction. Igiraneza and collaborators trained a model on HIV-1 envelope sequence embeddings using multi-task learning, where predictions are made across multiple antibody types simultaneously ^18^. In competitive tests, deep learning methods often improved prediction accuracy, especially for antibodies with limited training data. However, these models are also more difficult to interpret, making the identification of resistance mutations challenging.

In parallel work, Einav and collaborators have taken a different approach ^19–21^. They observed that the matrix of virus vs. antibody neutralization values exhibits a low-rank structure, with significant correlations across neutralization profiles. Einav et al. effectively learn a low-dimensional representation of the neutralization matrix to predict unobserved neutralization values. While this approach effectively captures antibody-virus similarities, it cannot predict resistance values for novel viruses because it relies on observed neutralization data rather than sequence or structural features.

A key challenge across these approaches is balancing predictive accuracy with generalizability. Sequence-based models risk overfitting due to their reliance on a large number of parameters, while low-rank matrix methods lack the ability to predict resistance patterns for novel viral sequences. Further advancements will likely require hybrid approaches that integrate both sequence-based and matrix-structure insights to enhance predictive robustness.

Here, we present a simple method for predicting viral sensitivity to antibodies based on finitely sampled neutralization data. We begin by quantifying similarities between antibodies based on their neutralization profiles across viral strains and clustering antibodies into distinct groups. Then, we project viral sequences into a low-dimensional space and train linear predictors of antibody sensitivity as a function of viral sequence features. Importantly, we use regularization to constrain the model such that the parameters for antibodies with similar neutralization profiles will also be similar. This allows us to leverage information across related antibodies to improve predictive power. Finally, we use a low-rank matrix approximation inspired by the work of Einav and collaborators ^19–21^ to further exploit antibody and virus correlations. Because of our focus on sharing information across groups of related antibodies and viruses, we refer to our approach as grouped neutralization learning (GNL).

We tested our approach using CATNAP, a large repository of HIV-1 virus-antibody neutralization data ^22^. Our method typically outperforms previous approaches that make the same types of predictions. GNL performs especially well for antibodies with limited observations and is notably more accurate for broadly neutralizing antibodies directed toward the variable loops and the CD4 binding site of the HIV-1 surface protein. Importantly, our method can successfully predict neutralization values for novel viruses without existing neutralization data, which we demonstrate in both CATNAP data and longitudinal data from a single donor. Overall, GNL improves our ability to predict virus-antibody neutralization values, which could be leveraged for practical applications such as antibody therapy design.

## Results

### Grouped neutralization learning framework

One of the principal challenges of predicting virus-antibody neutralization is sparse data. Neutralization experiments are generally low throughput, and the space of possible virus and antibody sequences is enormous. We assume that our data consists of a set of antibody-virus neutralization values and corresponding virus sequences. The neutralization values can be arranged into an *N*_*A*_ × *N*_*V*_ -dimensional matrix, *U*, where *N*_*A*_ is the number of antibodies and *N*_*V*_ the number of viruses, with the average neutralization value for antibody *µ* and virus *α* given by *u*_*µα*_ (for an example, see **Fig. 1a**). This value need not be defined for all antibody-virus pairs, and viruses may be included even if there are no neutralization values measured for them for any of the antibodies in the panel. Our goal is then to use the observed *u*_*µα*_ and virus sequences to predict the unobserved *u*_*µα*_. Here, we will assume that the neutralization values are quantified by half-maximal inhibitory concentrations (IC50s) on a logarithmic scale, but other metrics could also be used.

**Fig. 1.**
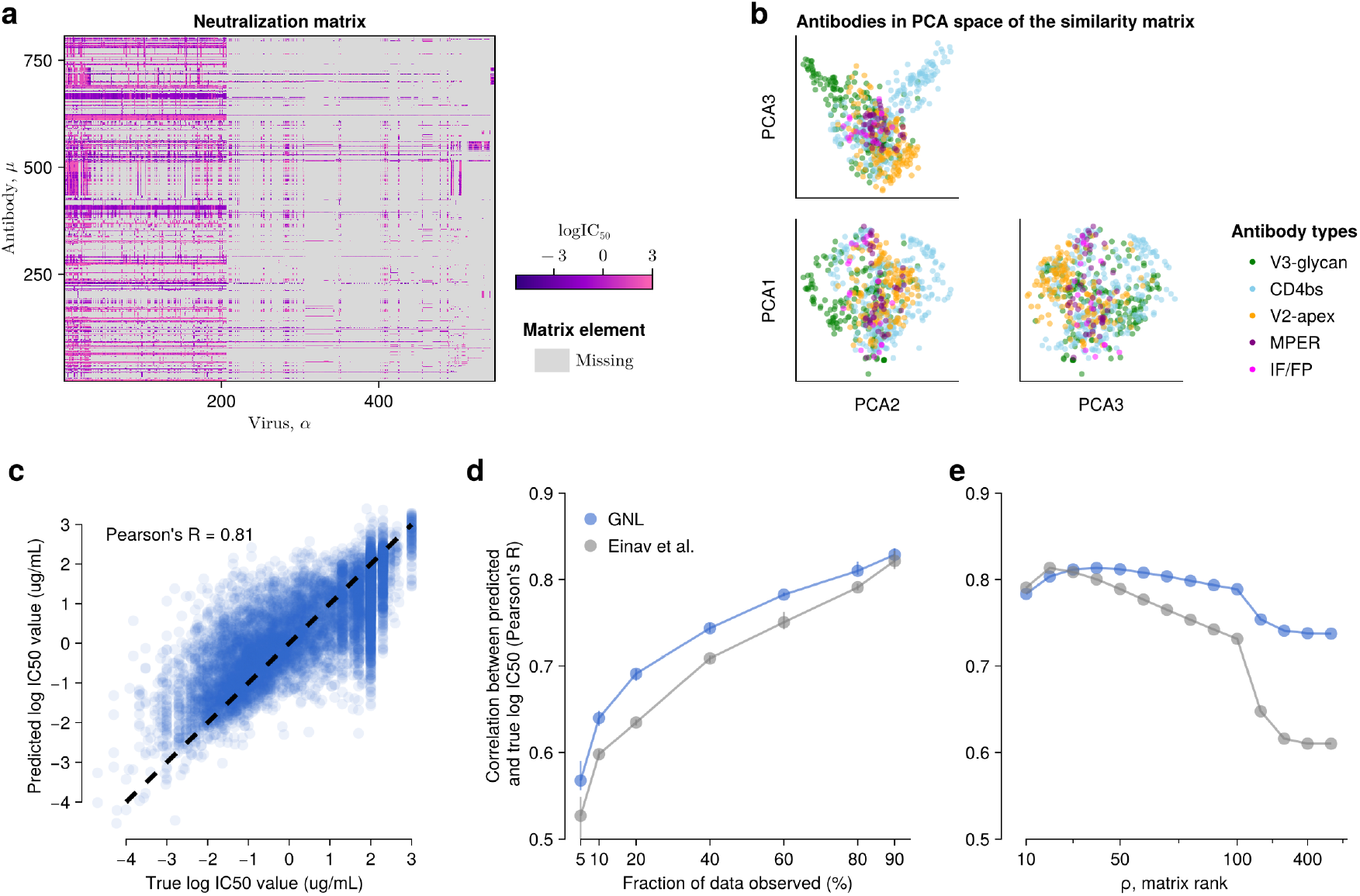
GNL improves virus-antibody neutralization predictions. **a**, Visualization of HIV-1 neutralization data retrieved from CATNAP ^22^. **b**, Antibody similarity visualized by projection onto the top principal components of the similarity matrix *w*_*µν*_. Antibodies are colored according to the part of the HIV-1 surface protein, Env, that they bind to. **c**, Comparison of neutralization values predicted by GNL with true, withheld values. **d**, For both GNL and the Einav et al. approach, predictive power (measured by Pearson’s *R* between true and predicted neutralization values) improves along with the amount of data used for training. **e**, Predictive power depends nonlinearly on the rank *ρ* used in the low-rank matrix approximation for both GNL and Einav et al. However, our approach exhibits stable and generally improved performance over a wide range of *ρ* values.

One of the main ideas of our approach is to leverage similarities between antibodies to improve predictive power. We also associate viral sequence features with antibody sensitivity or resistance, enabling predictions for novel sequences. Finally, we employ a low-rank approximation for the antibody-virus neutralization matrix, which makes use of common patterns in the matrix to refine the results. Below, we describe each of these steps (**Supplementary Fig. 1**) and their rationale.

### Grouping similar antibodies

To identify antibodies with similar neutralization profiles, we calculate a similarity matrix between all antibody pairs based on the observed neutralization values (see **Fig. 1b** for a visualization). The similarity 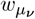 between antibodies *µ* and *ν* is defined as a product of the angular similarity of their neutralization vectors against the same viruses and the fraction of neutralization values that are shared between them (Methods). Thus, pairs of antibodies that have highly similar neutralization profiles against the same viruses, and those which have frequently been tested against the same viruses, will receive higher similarity scores.

We then apply a spectral clustering algorithm to group antibodies into a set of *K* clusters ^23^. Antibodies that exhibit similar neutralization activities across different viruses are grouped into the same groups. For the CATNAP data set, we used *K* = 100, though our results are robust to a range of values for *K* between 50 and 200. Most clusters contain one or a few antibodies, 10–20 clusters contain 10–20 antibodies, and only a few contain around 100 antibodies.

### Extracting sequence features and learning neutralization values

Here, we used a simple approach to embed viral sequence data into a lower-dimensional space. We computed the eigen-values and eigenvectors of the viral sequence covariance matrix and then projected each viral sequence onto the eigen-vectors corresponding to the *d* largest eigenvalues (Methods). Specifically, given a binary (one-hot) encoding of a viral sequence *α*, ***g***_*α*_∈ {0, 1} ^*L*^, we generate a projected sequence ***x***_*α*_ = Ξ***g***_*α*_. Here Ξ ∈ ℝ^*d×L*^ is a projection matrix, with *d* ≪ *L*. This approach is statistically efficient, and it takes advantage of correlated patterns of mutational variation observed in the data, which can affect virus sensitivity to antibodies. For example, in the broad HIV-1 CAT-NAP data set, we observed that different viral subtypes tend to occupy distinct regions in the eigenspace due to shared, subtype-defining mutations (**Supplementary Fig. 2**). While our main approach simply uses principal components, we also explored alternative dimensional reduction methods, including non-negative matrix factorization (NMF). However, the accuracy of our predictions changed little across different low-dimensional spaces as long as they provided statistically important features to distinguish viral sequences.

We used linear regression to learn the relationship between sequence features and neutralization values, inferring a separate family of regression models for each cluster of antibodies (Methods). For each cluster, we set *d* to be proportional to the number of unique viruses with measured neutralization values for antibodies in the cluster, up to a maximum of 100. We used the antibody similarity measures defined above, *w*_*µν*_, to penalize differences between the linear regression parameters ***θ***_*µ*_ and ***θ***_*ν*_ for each pair of antibodies *µ* and *ν* within the same cluster. In this way, similar regression parameters will be learned for antibodies with similar neutralization profiles (i.e., *w*_*µν*_ ∼ 1). In the limit that *w*_*µν*_ = 0, the regression parameters for antibodies *µ* and *ν* become independent. Overall, this approach allows the information from virus-antibody neutralization data to be shared among similar antibodies.

### Low-rank approximation for the neutralization matrix

Einav and collaborators found that typical virus-antibody neutralization matrices have a “low-rank” structure ^19–21^. In other words, the neutralization profiles for each virus and antibody are not completely unique; virus-antibody neutralization scores tend to be highly correlated across antibodies (for closely-related viruses) and across viruses (for antibodies that bind similar epitopes in a similar way). Thus, the neutralization matrix can be well-characterized by a number of modes much smaller than its naive dimensions. Our approach makes use of the same insight.

After inferring linear regression parameters for each antibody ***θ***_*µ*_, we use the regression model to predict the neutralization value *u*_*µα*_ for each virus-antibody pair that was not measured in the data set. We then refine this matrix by applying a low-rank approximation. Specifically, we used singular value decomposition (SVD) to factorize the preliminary *U* matrix, consisting of the measured *u*_*µα*_ with the unobserved entries filled by the regression model. We then selected the top *ρ* singular values and basis vectors to approximate the matrix. As in prior work ^19^, we selected the rank *ρ* to be the smallest rank such that the cumulative variance explained (equal to the sum of squared singular values) is greater than 95%. Our final predictions for the unobserved *u*_*µα*_ are then given by the entries of this low-rank approximation of the completed *U* matrix after regression.

In the past work, Einav et al. used robust principal component analysis or nuclear norm minimization to develop a low-rank approximation of the neutralization matrix ^19^. We also tested these methods, but in our analyses, we found that there was little difference between them and the simpler SVD factorization. Consistent with Einav et al., we also observed that the spectrum of singular values could reveal information about the neutralization matrix. We found that fraction of variance explained scales roughly as a power law for intermediate eigenmodes, ranging from around rank 20 to 100 (**Supplementary Fig. 3**).

### Neutralization and virus genetic data

In this study, we consider two types of neutralization matrices derived from monoclonal anti-HIV-1 antibodies and HIV-1 strains. The first is a neutralization matrix obtained from cross-sectional studies in the CATNAP database ^22^ provided by Los Alamos National Laboratory (LANL) ^24^. To ensure data quality, we first removed viruses and antibodies that had few associated neutralization values (*<* 10 entries for viruses, then antibodies that do not have any entries). We also removed viruses had not been sequenced.

After filtering, we obtained a data set with *N*_*A*_ = 806 antibodies and *N*_*V*_ = 1, 147 viruses, including 81,193 observed neutralization values. As indicated above, neutralization data in large data sets such as CATNAP are sparse: the measured neutralization values correspond to 4.5% of the total matrix entries.

The second data set comes from an HIV-1 intrahost longitudinal study, which tracked the development of broadly neutralizing antibodies in one donor, CH505 (ref. ^25^). We obtained virus-antibody neutralization values from Gao et al. ^26^ and corresponding HIV-1 Env sequences from LANL’s HIV sequence database ^24^. This matrix includes *N*_*A*_ = 22 antibodies and *N*_*V*_ = 109 virus sequences, with 1,723 measured neutralization values. In contrast to the CATNAP database, 71.9% of the possible neutralization matrix entries were measured in this data set. More details on the formatting of the neutralization matrices, the processing of genetic sequences, and the parameters of the models are provided in Methods and **Supplementary Table 1**.

### Overall GNL performance on CATNAP data

To test the performance of the GNL method, we masked a random fraction of virus-antibody neutralization values in the CATNAP data set. We then used GNL to predict the masked neutralization values and compared our predictions against the true measured values. First, we masked entries of *U* uniformly at random (i.e., each observed *u*_*µα*_ value was equally likely to be masked). **Figure 1c** shows a typical comparison between the true and predicted neutralization (log IC50) values when 20% of the data is masked, which are strongly correlated. In this example, there are 780 antibodies and 1,097 viruses in the withheld data.

We also compared our approach versus that of Einav et al. (ref. ^19^) as we varied the fraction of the data used for training and validation, and the rank *ρ* used in the low-rank matrix approximation. To measure typical performance, we repeated each test 10 times for each data fraction and *ρ* value. Overall, we observed that the correlation between true and predicted log IC50 values was typically higher for GNL than for the alternative (**Fig. 1d, e**). Comparatively, the results for GNL were especially better when the neutralization data were sparser. We also observed that GNL performance was more robust to changes in the rank *ρ*, with a wider plateau near peak performance. This robustness is especially important for practical applications, where the true neutralization values are unknown, because it implies that strong predictions can be obtained even without fine-tuning parameters.

### Determinants of prediction accuracy

Next, we aimed to identify conditions that make predicting virus-antibody neutralization values more or less difficult. We focused on two key factors in the neutralization data: the diversity of neutralization values and the amount of data available (**Fig. 2a-b**; see also **Supplementary Fig. 4**). Intuitively, antibodies with more observed neutralization values provide more information for training, potentially enabling higher accuracy. Antibodies with a broad range of neutralization values (i.e., ones with substantially different IC50 values for different virus strains) may also be more difficult to model than those with flatter neutralization profiles.

**Fig. 2.**
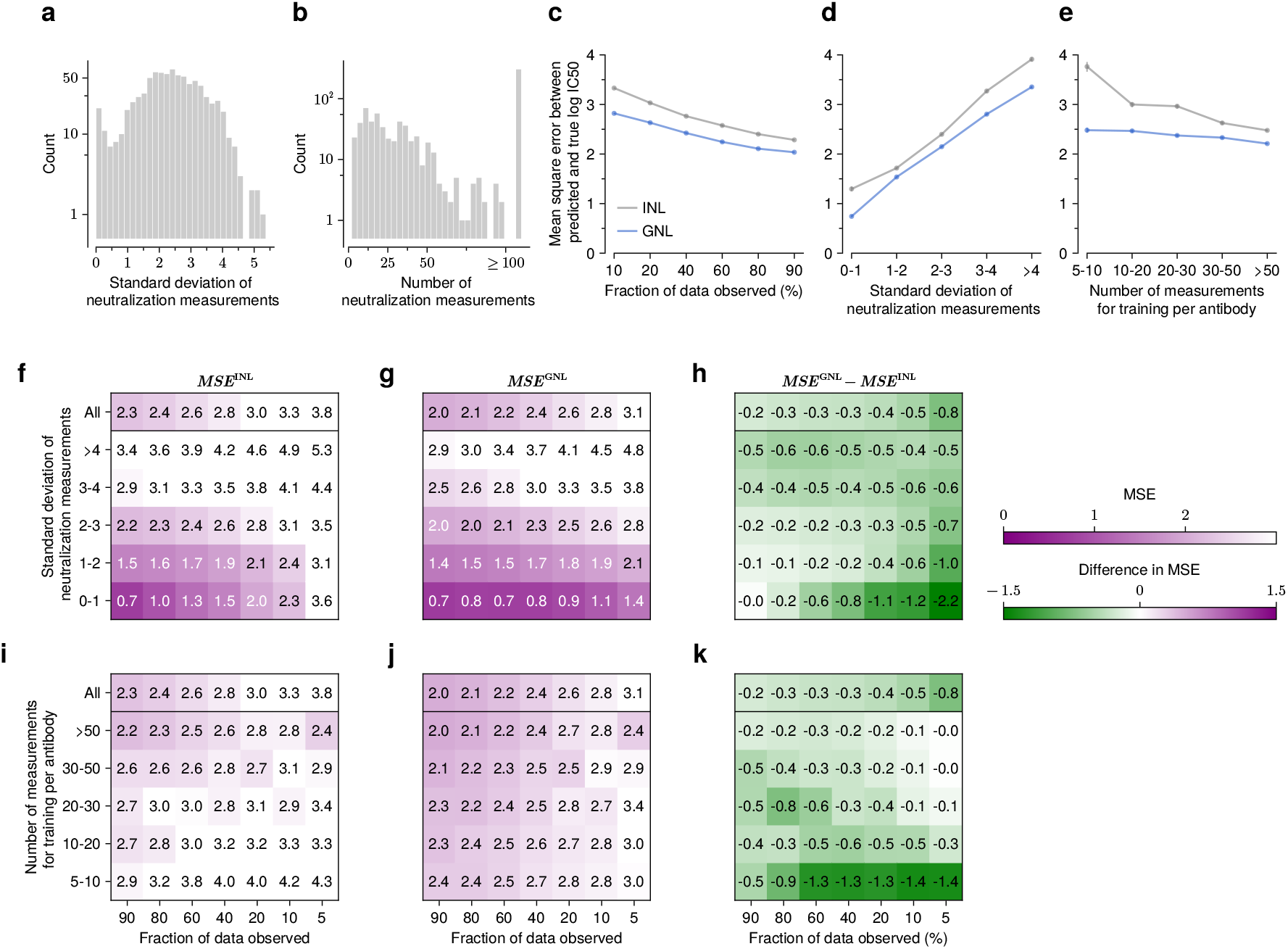
GNL improves predictions when neutralization measurements are widely divergent or when data is sparse. Overall distribution of the standard deviation of the neutralization measurements (**a**) per antibody, and their number of neutralization measurements (**b**). **c**, As the fraction of observed data decreases, accuracy drops (i.e., MSE increases); however, GNL consistently shows lower MSE, especially at smaller data fractions. GNL also performs better for antibodies with higher diversity, as measured by standard deviation (**d**), or with fewer observed measurements used for training (**e**). Greater diversity in neutralization measurements increases MSE across all withholding levels for both INL and GNL (**f-g**). However, GNL better mitigates this accuracy drop (**h**). Similarly, we observed that fewer training measurements lead to higher MSE across all withholding levels (**i-j**). However, the GNL method maintains lower MSE with fewer observations (**k**). **Supplementary Fig. 4** presents the same analysis including the Einav et al. method.

Because some subsets of the data contained only a small number of virus-antibody pairs, we used the mean squared error (MSE) between predicted and observed neutralization values to quantify accuracy in these comparisons. With few data points, correlation metrics such as Pearson’s *R* may not fully capture the correspondence between predictions and data. This is especially true for neutralization values with little variation, where predictions may be quite accurate on an absolute scale even if they are not strictly ordered correctly. For example, antibodies VRC34-UCA, VRC34.05, vFP5.01, A12V163-a.02, DH650.8, and m66 have more than 200 neutralization observations, but the variance in measured log IC50 values for each is less than 0.1.

For reference, we compared our results against a simple version of our model where we treat each antibody independently. This independent neutralization learning (INL) model is equivalent to simple linear regression to predict neutralization values for each antibody. Overall, we found that MSE values tend to decrease as the amount of training data increases and as the variance in the training data decreases (**Fig. 2c-e**). Errors for GNL were always smaller than those for INL, demonstrating the value of shared information across antibodies with similar neutralization profiles.

We grouped antibodies into five classes according to the standard deviations of their neutralization values. **Figure 2f-g** shows that MSE increases as the diversity of log IC50 values increases. Consistent with analyses in **Figure 1**, we set the fraction of data observed as 80%. More generally, when large fractions of the observed data are used for training, the difference in MSE values between the GNL and INL methods becomes more pronounced when the diversity of neutralization measurements is higher (**Figure 2h**), a setting in which learning neutralization values is more challenging.

We further partitioned antibodies based on the number of neutralization measurements used for training (**Fig. 2i-k**). Naturally, MSE values tend to be larger for antibodies with fewer training data. However, the advantage of GNL over INL is especially prominent in the cases where data is most restricted, including a small fraction of the data observed, smaller training data sets, and less variation in the training data (**Fig. 2h, k**). This suggests that the grouping mechanism in GNL, which allows predictive information to be shared across similar antibodies, is especially helpful in learning genotype-to-neutralization mappings with sparse data.

### Comparison against alternative approaches for virus-antibody neutralization prediction

Alongside low-rank approximation methods, there are other families of neutralization prediction approaches, including ones based on machine learning ^10–12,15,17^. One widely used tool is SLAPNAP (Super Learner Prediction of NAb Panels) ^12^. SLAPNAP has been applied to several studies and predicts neutralization values for single and combinations of bnAbs using viral envelope protein sequence features ^13,14^. The underlying models include random forests and boosted regression trees with *L*_1_ regularization. For combinations of bnAbs, SLAPNAP uses the Bliss-Hill model ^27^, which estimates single bnAb neutralization curves via a Hill function and combines their effects based on Bliss independence.

To compare SLAPNAP’s performance with the GNL method as well as the Einav et al. method ^19^, we first replicated their training settings. In particular, we trained each model using *V* -fold cross-validation. In this setup, the data is divided into *V* parts: *V* − 1 are used for training, and the remaining part is used for validation. This is repeated *V* times. We set *V* = 5, following prior work ^12^. To ensure a fair comparison, we used the same neutralization data set (from CATNAP, as of July 23, 2021) previously tested for SLAPNAP. The selection and grouping of antibodies for this validation were also based on prior work ^12^. Compared to the full CATNAP data set, the selected antibodies often had a large number of neutralization measurements (>200, for comparison see **Figure 1b**).

Figure 3. reports CV-R values for GNL, SLAPNAP, and Einav et al. methods on this data set, showing that GNL performs well across a wide range of antibodies. Across all antibodies, GNL achieved an average CV-R of 0.60, compared to 0.51 for SLAPNAP and 0.43 for the Einav et al. approach (**Supplementary Table 2**). For antibodies DH270.5, DH270.6, and VRC38.01, SLAPNAP was unable to produce a CV-R value. These antibodies have fewer neutralization measurements than other antibodies in this set, and thus the lack of a CV-R result may be due to a data restriction in SLAPNAP. To ensure a fair comparison across methods, we thus excluded antibodies for which SLAPNAP did not produce CV-R values.

**Fig. 3.**
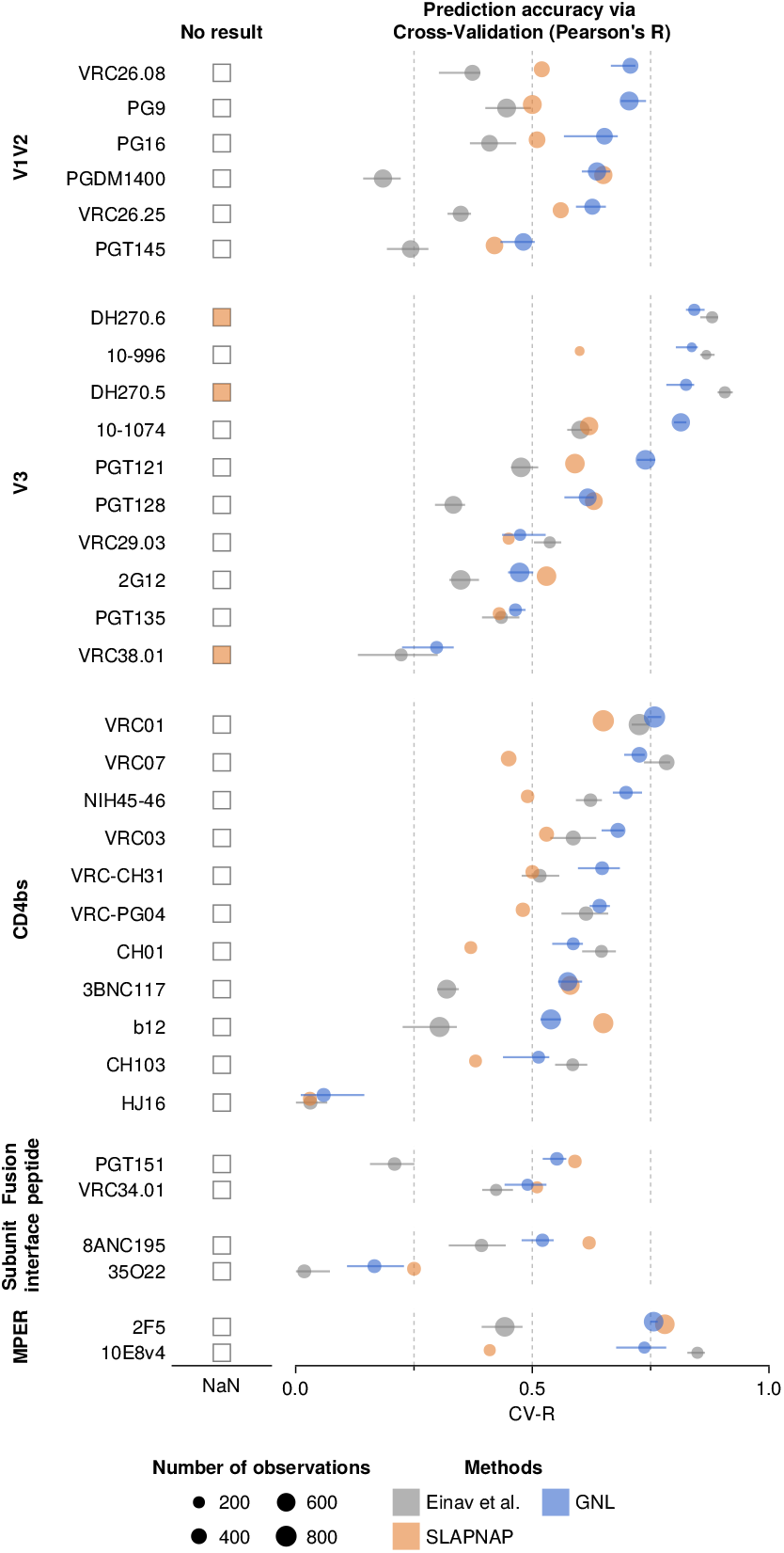
Performance comparison of GNL and existing methods on CATNAP data. Cross-validation Pearson’s *R* (CV-R) for neutralization predictions across a spectrum of antibodies with GNL, SLAPNAP ^12^, and Einav et al. For a few antibodies (DH270.5/6 and VRC38.01), SLAPNAP produced no output. The average CV-R of GNL across all antibodies (0.60) exceeded SLAPNAP (0.51) and Einav et al. (0.46) (**Supplementary Table. 2**). GNL performance was especially strong for antibodies binding the variable loops (V1/V2 and V3) and the CD4 binding site.

### Zero-shot prediction of neutralization values for novel viruses

Next, we tested the ability of GNL to perform “zero-shot” predictions of antibody-virus neutralization values for viruses that never appeared in training data. Predicting neutralization values for novel viruses is challenging because it requires learning generalizable relationships between virus properties (e.g., sequence, structure) and resistance, rather than simply relying on observed neutralization patterns. Matrix completion methods that depend exclusively on observed virus-antibody pairs cannot make predictions for entirely new viral sequences. To evaluate GNL’s zero-shot capabilities, we considered two experimental scenarios using longitudinal data from a single HIV-1 infected donor, CH505, and cross-data set predictions between independent neutralization studies.

As a first example, we considered longitudinal neutralization data from donor CH505, who was monitored over five years. The CH505 data set includes neutralization values for *N*_*A*_ = 22 antibodies tested against *N*_*V*_ = 109 viruses over six time points ^25,26^. To validate our approach, we first confirmed that GNL could accurately predict randomly withheld neutralization values, achieving strong correlation (*R* = 0.89) when 80% of the data was observed (**Supplementary Fig. 5b**, see also Methods for the rank optimization scheme). We then examined two temporal prediction scenarios: predicting neutralization values for past viral sequences using models trained on later time points, and predicting future viral sequences using models trained on earlier data.

For temporal predictions, we split the neutralization data based on collection time, using early weeks post-infection (28, 96, and 140 weeks) and later weeks (210, 371, and 546 weeks) as separate training and validation sets. When predicting antibody-virus neutralization values for early viruses based on late infection data, GNL achieved Pearson’s *R*-values of approximately 0.8 across all time points (**Supplementary Fig. 4c**). For the more challenging task of predicting neutralization values for future viral sequences based on prior (early infection) data, GNL maintained good predictive accuracy (Pearson’s *R* ranging from 0.7 to 0.55) despite the continued accumulation of mutations (**Fig. 4b**). This reflects the ability of GNL to learn generalizable maps from viral sequence to neutralization.

**Fig. 4.**
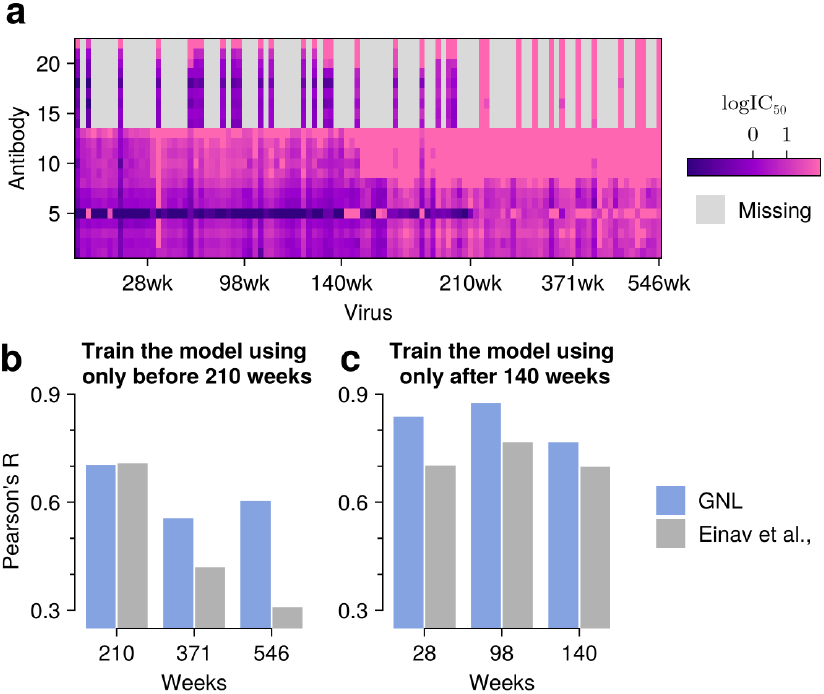
GNL accurately predicts neutralization values for viral sequences across evolutionary time. **a**, Visualization of the neutralization matrix, consisting of *N*_*A*_ = 22 antibodies and over *N*_*V*_ = 109 viral sequences sampled across six time points. **b**, Prediction of the neutralization values for past viral sequences. Neutralization values were predicted for sequences sampled at or before 140 weeks, using a model trained only on data collected after 140 weeks. **c**, Prediction of the neutralization activities for future viral sequences. The model was trained on values observed up to 140 weeks, and predictions were tested on sequences sampled after 210 weeks. GNL maintains accuracy even as viral sequences diverge from training data over time.

In a second test, we evaluated whether GNL could predict neutralization values for an individual host using only data from the broader CATNAP database. We trained models exclusively on CATNAP data and predicted neutralization values for the CH505 individual host data set, focusing on the two antibodies that are present in both the CATNAP and CH505 data sets (**Fig. 5a**). Since the CATNAP database and individual host data are largely disjoint, with none of the CH505 virus sequences present in CATNAP, this represents a stringent test of zero-shot predictive power. Here, GNL achieved meaningful predictive accuracy with a Pearson’s correlation of *R* = 0.55 between predicted and true neutralization values (**Fig. 5c**). In comparison, predictions based on matrix completion are highly limited due to the absence of the specific test virus sequences in the training data (**Fig. 5b**). These results show that GNL can successfully transfer neutralization patterns from cross-sectional studies to predict virus-antibody interactions in independent data sets.

**Fig. 5.**
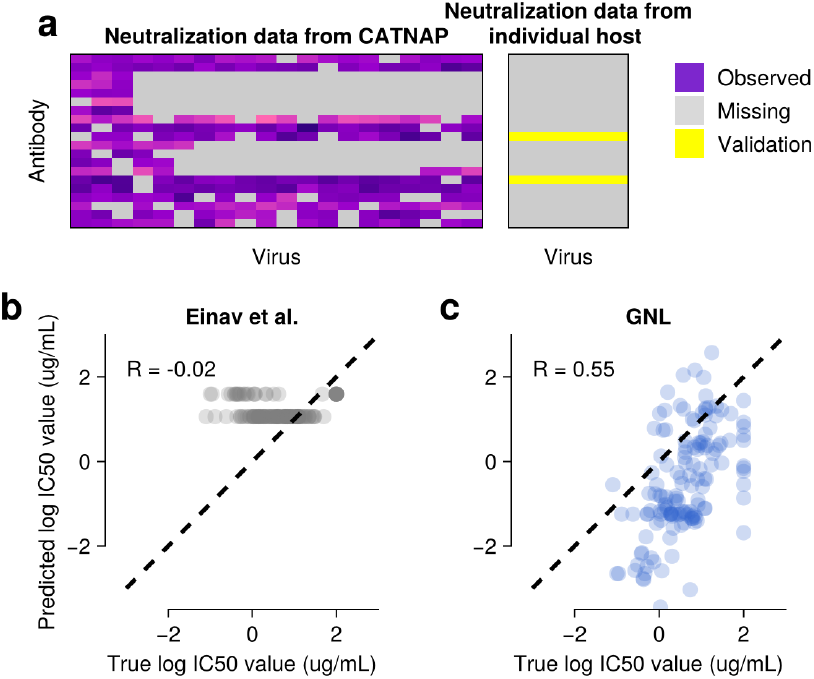
Neutralization values can be predicted for novel viruses using cross-data set transfer learning. **a**, Schematic showing a subset of the CATNAP database (left) and the individual host (CH505) neutralization matrix to be predicted (right). CATNAP contains *N*_*A*_ = 806 antibodies and *N*_*V*_ = 1, 147 viruses, while the CH505 matrix includes 22 antibodies and 109 viral strains. Only two antibodies are shared between the data sets. **b**, Comparison of true neutralization values from CH505 and predicted values using the Einav et al. method ^19^, which can only predict mean values from the training data. **c**, GNL achieves meaningful predictive accuracy for the same task (Pearson’s *R* = 0.55, Spearman’s *ρ* = 0.6, and linear regression slope 1.2).

## Discussion

Quantifying the efficiency of drugs or antibodies against pathogens is a fundamental task in biology. However, experimental neutralization measurements can be resource-intensive, and evaluating all possible combinations of treatments and pathogens is infeasible. We developed grouped neutralization learning (GNL), a simple method for predicting viral sensitivity to antibodies based on finitely sampled neutralization data. GNL employs three key ideas to boost predictive power: sharing information among antibodies with similar neutralization profiles, mapping from viral sequence to neutralization using regularized linear regression, and refining predictions using a low-rank matrix approximation, inspired by refs. ^19,21^. In contrast with previous approaches based on matrix completion, GNL can use sequence features to predict neutralization values even for viruses with no prior neutralization data. We used GNL to predict half-maximum inhibitory concentration (IC50) values, but in principle the same approach could be applied to other similar metrics.

In competitive tests, our method compares favorably with the current state-of-the-art across a variety of scenarios. Our approach particularly excels in challenging conditions where data is sparse, using shared information from antibodies with similar neutralization profiles to enhance predictions (**Fig. 2**). GNL is also especially robust to the choice of the matrix rank used in the low-rank matrix approximation step (**Fig. 1**). This robustness is particularly valuable for practical applications where the rank must be chosen heuristically.

Our ability to predict neutralization values for novel viral sequences enables several important biological applications. In a test on longitudinal data from a single donor, GNL could predict neutralization patterns for future viral sequences based on prior sequence and neutralization data (**Fig. 4**). GNL was also capable of “transfer learning,” using antibody-virus neutralization data from the CATNAP database to predict neutralization values for viruses in the CH505 data set, despite the absence of these sequences in CATNAP data (**Fig. 5**). These capabilities could be used to inform antibody-based treatments and to aid in understanding virus evolution to escape antibodies ^25,26,28–31^.

Our approach addresses several important challenges in neutralization prediction. GNL effectively learns from large but sparse neutralization data sets such as CATNAP, where neutralization values for only around 4.5% of all antibody-virus pairs are measured. As data sets grow larger, they are also almost certain to increase in sparsity. Sharing information among antibodies with similar neutralization profiles is especially beneficial for antibodies with limited training data. Partitioning antibodies into groups also improves the computational efficiency of GNL. Combining sequence-based predictors with low-rank matrix refinement allows us to incorporate both specific sequence features associated with antibody sensitivity or resistance and general patterns of neutralization across antibodies.

There are also several limitations of GNL that should be considered. First, we assume a linear relationship between viral sequence features and (log) neutralization values, which may not capture more complex nonlinear mutation effects. Similarly, our dimensional reduction approach uses principal components of viral sequence covariance, which may not capture the most relevant features for predicting neutralization. However, this is not an inherent limitation of GNL, which could be adapted to use arbitrary sequence features for prediction. Finally, the similarity metric that we use for grouping antibodies depends on overlapping neutralization measurements, which may be more limited for newly discovered antibodies or those tested against different panels of viruses.

GNL could be extended in multiple ways in future work. Our framework could apply the Bliss-Hill approach to estimate neutralization values for combinations of antibodies, provided that concentration-dependent neutralization values are available for multiple viruses and antibodies ^27,32^. We could further extend our model to consider more complex relationships between viral genotype and neutralization (for example, ones including epistasis ^33–37^), which could improve predictive power. The interpretability of sequence features could be enhanced by incorporating biologically meaningful representations such as sequence motifs ^38,39^ or known resistance mutations, rather than relying solely on principal component analyses. While we have focused on HIV-1 in this work, our framework could readily be extended to other rapidly evolving pathogens where neutralization data and genetic sequences are available.

## ACKNOWLEDGEMENTS

This work of K.S.S., J.P.B., and R.M.L. was supported by the National Institute of Allergy and Infectious Diseases of the NIH award number R01AG123456.

## AUTHOR CONTRIBUTIONS

All authors contributed to the design of the study, interpretation of the results, and writing of the article. K.S.S. performed simulations and computational and mathematical analyses. J.P.B. supervised the project.

## Methods

### Dimensional-reduction of viral sequences

To transform genetic sequences into an amenable data structure, we first process them into one-hot encoded sequences. For example, consider an aligned amino acid sequence 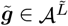, where 𝒜={A, C, …, Y, -} is the set of amino acids, including the alignment gap symbol ‘-’, and 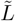 is the length of the amino acid sequence. To convert this amino acid sequence 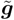 into a numerical form, we encode each amino acid as a *q* = 21-dimensional vector (corresponding to all possible amino acids and the gap symbol), consisting of zeros except for a single one at the position encoding the amino acid. By concatenating these *q*-dimensional column vectors 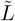 times, we obtain the one-hot encoded sequence, which consists of 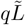 elements. Since the vast majority of elements in the one-hot encoded sequence are conserved across all considered viral sequences, we discard these conserved sites and retain only the variable sites. This results in ***g*** ∈ {0, 1}^*L*^, a one-hot encoded sequence that contains only the variable sites, where *L* represents the number of variable sites in the one-hot sequence ensemble. *L*, the number of variable sites of one-hot encoded sequences (approximately 7,000), is much larger than the number of neutralization values available for most antibodies (ranging from 2 to about 1,000). To learn a meaningful model while avoiding overfitting, we first project the one-hot encoded sequences into a lower-dimensional feature space. Formally, we consider a linear transformation of the one-hot encoded sequence:

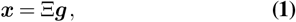

where Ξ ∈ ℝ^*d×L*^ is a projection matrix that maps the one-hot encoded sequences to the feature space. Each row of Ξ corresponds to a single feature and ***x*** represents the projected sequence (i.e., consisting of the *d* features of the single viral sequence), which will then be used to predict neutralization values.

In this study, we used the eigenmodes of the covariance matrix of genetic sequences as the Ξ matrix. Specifically, we obtained the covariance matrix as *C* = ⟨ ***gg***⟩^⊤^− ⟨ ***g***⟩ ⟨ ***g***⟩ ^⊤^, where ⟨·⟩ denotes the average over the viral genetic sequences in the data set. We then computed the eigen-vectors ***ξ***_*i*_ and eigenvalues *λ*_*i*_ of *C*. One can then write 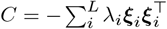, where ***ξ***_*i*_ and *λ*_*i*_ are the *i*-th eigenvector and eigenvalue, respectively, with *λ*_*i*_ ≤ *λ*_*i*+1_ ≤ 0 for *i* ∈ {1,…, *L* − 1}. Finally, we constructed the Ξ matrix by selecting the first *d* eigenvectors as the row vectors of Ξ, such that the *j*-th element of *i*-th eigenvector Ξ_*ij*_ = (***ξ***_*i*_)_*j*_.

To learn the neutralization values from the projected sequences ***x***, we set *d* based on the number of observed neutralization values. Specifically, we define |𝒩_*µ*_ | as the number of observed neutralization values and set:

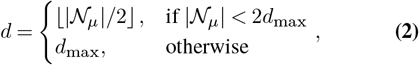

where ⌊*a*⌋ represents the greatest integer that is equal to or smaller than *a*, and *d*_max_ denotes the maximum dimension of the feature vector, which we set as *d*_max_ = 100 throughout. As we will discuss later, in the case of the grouped learning method, we replace 𝒩_*µ*_ with 𝒩_*k*_, the union of all antibodies 𝒩_*µ*_, where *µ* belongs to the*k*-th antibody group, such that 𝒩_*k*_ = ⋃_*µ*∈*k*-th antibody group_ 𝒩_*µ*_ The choice of the Ξ matrix, which defines the low-dimensional space, is flexible. As long as the proposed low-dimensional space effectively captures key features of the virus sequence ensemble, any appropriate matrix can be used. For example, non-negative matrix factorization (NMF) or other low-dimensional spaces that learn interpretable features such as structural or evolutionary signatures ^31–33^, could be applied for this purpose.

### Neutralization similarities and grouping of antibodies

We consider the following antibody similarity matrix based on partially observed neutralization measurements. Let 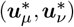 be a pair of neutralization vectors of antibodies *µ* and *ν* that are restricted to the *n*_*µν*_ viruses where both antibodies have neutralization values (i.e., the intersection, 𝒩_*µ*_ ∩ 𝒩_*ν*_). The similarity matrix is then defined as

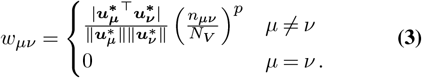

Here, | · | denotes the absolute value and ∥ · ∥ represents the *L*_2_-norm (e.g., for a vector ***v*** with elements 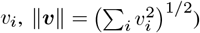. The first term represents the cosine or angular similarity between the two vectors in the restricted space, which ranges between zero and one. The second term ensures that the similarity weight increases along with *n*_*µν*_, the number of viruses that both antibodies have been tested against. The exponent 0 ≤ *p* controls the extent of this tuning. When *p* = 0, the fraction of overlap between the two antibodies is unimportant and this term is simply equal to one for all pairs of antibodies that have any overlap. As *p* becomes larger, the emphasis on the overlap increases. In our study, we set *p* = 0.2 but the results were robust for *p* ∈ [0.1, 0.4]. If one uses different neutralization data, the *p* value may need to be adapted so that the neutralization profiles are similar within the same group of antibodies.

As we will discuss in the next section, the similarity matrix ensures correlations between models that predict the neutralization activity of similar antibodies. The low-dimensional representation of the antibodies is also derived from the eigenmodes of the similarity matrix (**Fig. 1b**. Antibodies are grouped based on the similarity matrix, which is used to compute the graph Laplacian matrix for applying the spectral clustering algorithm ^34^.

### Learning neutralization values

To predict the missing neutralization values, we learn the relationship between sequences and neutralization values. To do this, we define a model that reproduces the neutralization value as *u*_*µα*_ = *f* (***x***_*α*_ |***θ***_*µ*_)+*ϵ*_*µα*_. Here, *f* : ℝ^*d*^ → ℝ is a function, ***θ***_*µ*_ is a model parameter that captures the neutralization values of antibody *µ*, and *ϵ*_*µα*_ is the uncertainty, which we assume is normally distributed with a mean of zero. Let ***θ*** and ***x*** be 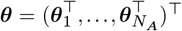 and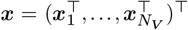, respectively. Using Bayesian rule, the optimal model parameters are obtained as a maximum a posterior estimator, given by

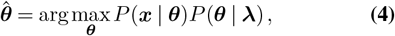

where ***λ*** = (*λ, λ*^*′*^)^⊤^ is a set of parameters characterizing the distribution of ***θ***, which we will discuss below.

Here, the likelihood function is given as

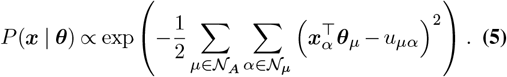

For the prior distribution, we assume that antibodies with similar neutralization profiles share similar model parameters. Formally, the standard deviation of the difference between these parameters is inversely correlated with the similarity weights, therefore we represent it as

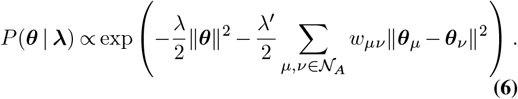

The optimal parameter, derived from the logarithm of the maximization of a posterior distribution, is the solution to its stationary condition, expressed as

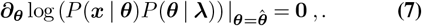

To solve the stationary equation, let us define a matrix as

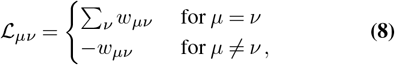

which is equivalent to the weighted graph Laplacian matrix ^34^, and its lifted matrix L as

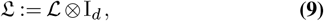

where ⊗ and I_*d*_ represent the outer product and *d*-dimensional identity matrix, respectively. Here, 𝔏 is a block diagonal matrix, with entries 𝔏_*µν*_I_*d*_ in the block spanning row indices from (*µ* − 1)*N*_*A*_ + 1 to *µN*_*A*_ and column indices from (*ν* − 1)*N*_*A*_ + 1 to *νN*_*A*_. Finally, the analytical expression of the optimal model parameter 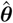 can be expressed as

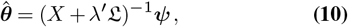

with

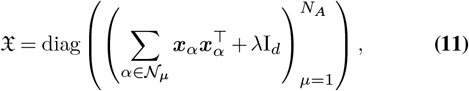

which is a block diagonal matrix consisting of *N*_*A*_ *d*-by-*d* blocks, and

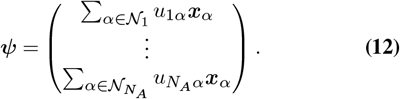

In this construction, and obtained equations (Eq. (10) to Eq. (12)), when the ℒ matrix is diagonal (ℒ_*µν*_ = 0 for *µ* ≠ *ν*), that is, all the similarity weights are zero, resulting in the 𝔏 matrix becoming a trivial (block) diagonal matrix. In this case, Eq. (10) is independent for each antibody and recovers the naive linear regression solution. We refer to this as independent neutralization learning (INL) method. However, as long as the elements of ℒ_*µν*_ ≠ 0, the equations for antibodies *µ* and *ν* become dependent, and their influences are interconnected. In particular, a group of antibodies with similar higher weight values influences each other, and we refer to this as the grouped neutralization learning (GNL) method.

### Efficient computation for grouped learning methods

As discussed above, the similarity weight can be used to identify distinct antibody groups by applying a standard spectral clustering algorithm ^34^. Once antibodies are grouped by *K* groups (where *K* is a positive integer and a hyperparameter) based on similar neutralization profiles, the equation to be solved Eq. (10) is reduced to smaller sub-problems: assuming that the weight matrix involves *K* disjoint blocks, then the full Laplacian matrix is divided into *K* sub-Laplacian matrices, 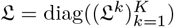, where 𝔏^*k*^ is *k*-th sub-Laplacian matrix. Therefore, the original problem Eq. (10) is reduced to solve the following sub-problem for each *k*:

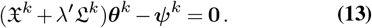

Here 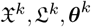, and ***ψ***^*k*^ are ones that associated with the *k*-th group.

This divided rule efficiently reduces computational complexity, especially when similar antibodies are classified into smaller groups. This divided rule allows us to employ adaptive group-dependent regularization strength 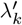, and group-dependent dimension *d*_*k*_, which is given by *d*_*k*_ = ⌊|𝒩_*k*_ | ⌋*/*2 if |𝒩_*k*_ | *<* 2*d*_max_ other wise *d*_*k*_ = *d*_max_, where 𝒩_*k*_ represents the union of 𝒩_*µ*_ for all antibodies *µ* belong to the *k*-th antibody group.

### Iterative matrix inversion algorithm

The matrices 𝔏 and 𝔛 are individually invertible with low computational costs, as 𝔏^−1^ = ℒ^−1^ ⊗ I_*d*_, and the inversion of a block diagonal matrix is computationally simpler. However, combining 𝔏 and 𝔛 disrupts these advantageous properties. Therefore, inverting the combination of them could take a longer time. To accelerate the process, we could employ an iterative algorithm to obtain the solution.

The Laplacian matrix ℒ should have at least one singular mode, as ℒ**1** = **0**, with **1** is the *d*-dimensional vector whose elements are all 1. If antibodies are divided into *K* distinct and disjoint classes, then the rank of the Laplacian matrix is *r* = *N*_*A*_ − *K*. We denote the Laplacian matrix as 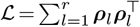, where ***ρ***_*l*_ is the *l*-th eigenvector corresponding to strictly positive eigenvalues. The extended Laplacian matrix 𝔏 is then represented as follows:

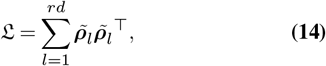

With

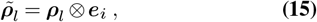

where ***e***_*i*_ has one at the *i*-th element and all other elements are zero. Based on the efficient computation algorithm for obtaining the inverse of the sum of matrices ^35^, the inverse of the matrices can be solved iteratively by:

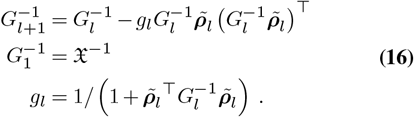

By iteratively solving 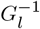 up to *l* = *rd* times, we get 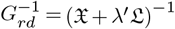.

The naive matrix-vector multiplication for 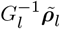 requires 𝒪 (*r*^2^*d*^2^) operations; however 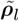 is a sparse matrix and it requires only 𝒪 (*r*^2^) operations, the outer-product also takes solely 𝒪 (*r*^2^) operations, and requires updates only 𝒪 (*r*^2^) elements at each *l*-th iteration. Therefore, after repeating the operation 𝒪 (*rd*) times, the total computational cost is only 𝒪 (*r*^3^*d*), which is much smaller than 𝒪 (*r*^3^*d*^3^) of the naïve inverse of the matrix.

### Processing the neutralization data

We retrieved IC50 neutralization values from the CATNAP database provided by HIV sequence database at Los Alamos National Laboratory (LANL) ^36^. By November 2024, over 172,000 IC50 neutralization values were available, and more than 159,000 of these values were tested with one of 806 monoclonal antibodies and 2,486 viral strains. Below, we describe the data processing steps, which are largely consistent with those used in Einav et al.

Some IC50 values exceeded experimental thresholds, typically above 50 or below 0.01. To address this, we replaced values above the threshold with double the threshold and those below it with half the threshold (e.g., values >50 were set to 100, and values <0.01 were set to 0.005). After this adjustment, we transformed the raw IC50 values using the logarithm. This transformation aligned the scale with viral fitness under antibody pressure and emphasized smaller neutralization values (same as the prior work ^37^). We then identified unique antibodies and virus strains, arranging the neutralization values into a matrix with rows corresponding to antibodies and columns corresponding to viruses. For antibody-virus pairs with multiple neutralization values, we computed the arithmetic average of the (log-transformed) values. All sources of experimental data were treated equally (same as ref. ^37^). Although we haven’t applied it due to the limited data, normalization of neutralization values could potentially reduce bias between experiments if sufficient values for certain antibody-virus pairs are available across multiple experiments. At this stage, we obtain *N*_*A*_ = 806 antibodies and 2,486 viruses, resulting in 90,738 neutralization values (about 4.5% of the total matrix.) We limited our analysis to virus strains with available envelope protein sequences, ultimately selecting 2,229 strains. Additionally, to improve data quality, we excluded virus strains with fewer than 10 neutralization values, resulting in *N*_*V*_ = 1, 147 strains. After filtering out antibodies without neutralization values (none of the antibodies were the case, so *N*_*A*_ = 806), we were left with 81,193 neutralization values, resulting 4.5% of the filtered neutralization matrix.

For the intrahost HIV neutralization data, we retrieved the neutralization values from Gao et al. ^38^. As noted in the main section, the neutralization matrix consists of *N*_*A*_ = 22 antibodies, *N*_*V*_ = 109 viruses, and a total of 1,723 neutralization values.

### Processing the viral sequences

In this study, we focused on the Env gene, which encodes the virus envelope protein targeted by antibodies. Using the alignment software provided by LANL ^39^, we aligned the viral Env protein sequences to the HIV reference strain HXB2. This allowed us to map amino acid positions to standardized indices commonly used in HIV research. To simplify analysis, we only considered positions that were not alignment gaps in the HXB2 sequence. This resulted in 856 positions that were present in most viral sequences. We converted the aligned viral Env sequences into one-hot encoded vectors. Each position (21 possibilities: 20 amino acids plus a gap) was represented as a binary vector with a single ones corresponding to the observed allele and zero elsewhere. This produced 856 × 21 = 17, 976 binary features per sequence. Since many positions were conserved across sequences, we retained only variable positions, yielding *L* = 7, 802 variable positions. The resulting one-hot encoded sequences still contain a large number of variables. Further dimensional reduction is described in the following section.

### Details of conditions for genotype-neutralization learning

The number of principal components used depended on the number of observed neutralization values. We set the minimum number of components to 2 and the maximum to 100, choosing half the number of observations for most cases. The strength of the *L*_2_ regularization on the parameter was fixed at *λ* = 5, and the regularization for the similarity of the model was fixed at *λ*^*′*^ = 1 for all simulations.

### Low-rank approximation

As detailed in the main text, we used standard singular value decomposition (SVD) instead of the robust principal component analysis (rPCA) method employed in previous studies by Einav et al. ^37^. Our results showed no significant differences between SVD and rPCA.

Before applying the low-rank approximation, missing values were completed using the GNL model based on viral sequences. For the Einav et al. method, missing values were replaced by mean estimates calculated from observed data for the corresponding antibodies and viruses. Mean values were subtracted before performing low-rank approximation and added back afterward, following procedures from the Einav et al. study ^37^.

The optimal rank value is determined as the minimum rank value where the variance explanation exceeds 95%, as per the optimization scheme of the Einav et al. method ^37^. The variance explanation for the matrix rank *ρ* is defined as the cumulative sum of the squared eigenvalues, 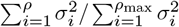. Here, *σ*_*i*_ is the *i*-th singular value (*σ*_*i*_ ≤ *σ*_*i*+1_), and *ρ*_max_ is the total number of singular modes.

The accuracy of neutralization imputation depends on the matrix rank. In **Fig. 1e** and **Supplementary Fig. 3**, we show how accuracy varies with matrix rank *ρ*: the accuracy changes significantly with rank in the method by Einav et al., while the GNL method is more robust and generally achieves higher accuracy. **Supplementary Fig. 1** also compares the variance explanation profiles and highlights the optimal rank values.

## Code availability

Sets of data and computer codes available in the GitHub repository: https://github.com/bartonlab/paper-ic50-prediction. The repository contains neutralization datasets and HIV-1 envelop genetic sequences. The most recent neutralization data and HIV viral sequences datasets are obtained from the CAT-NAP (http://hiv.lanl.gov/catnap) and LANL (https://www.hiv.lanl.gov) databases, respectively.

---

**Supplementary Fig. 1.**
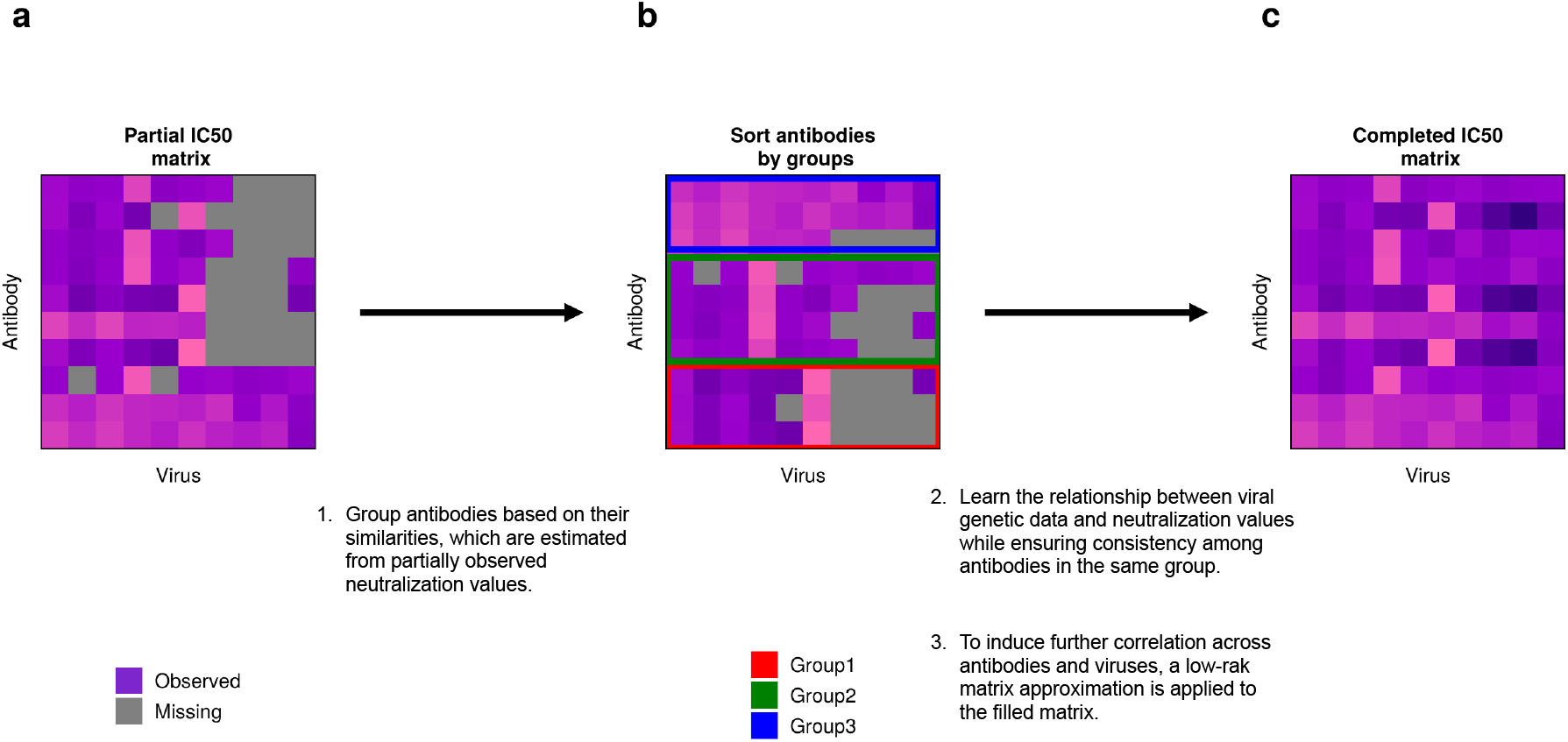
Overview of the neutralization imputation processes. **a**. The partially observed neutralization matrix (columns: viruses, rows: antibodies) is used as input to impute the missing elements (gray scale) of the neutralization matrix. **b**. Schematic representation of grouping antibodies based on the similarity of their neutralization values. Within the same group, neutralization activities are more similar than those between different groups. **c**. Temporarily complete missing values using models that learn the relationship betwee}n viral genetic data and neutralization values. We employ the novel grouped learning method, training the model to simultaneously learn the relationships between neutralization values and viral genetic sequences for a group of antibodies that share similar neutralization profiles (Methods). Here, to effectively reduce the number of model parameters and avoid the common over-parametrization issue, we applied a standard dimensional reduction technique to viral sequences and trained the model using the projected sequences. **d**. Obtain the complete neutralization matrix, derived from the low-rank matrix approximation of the temporarily completed matrix from step **c**. The optimal matrix rank was determined based on the distribution of eigenvalues (Methods).

**Supplementary Fig. 2.**
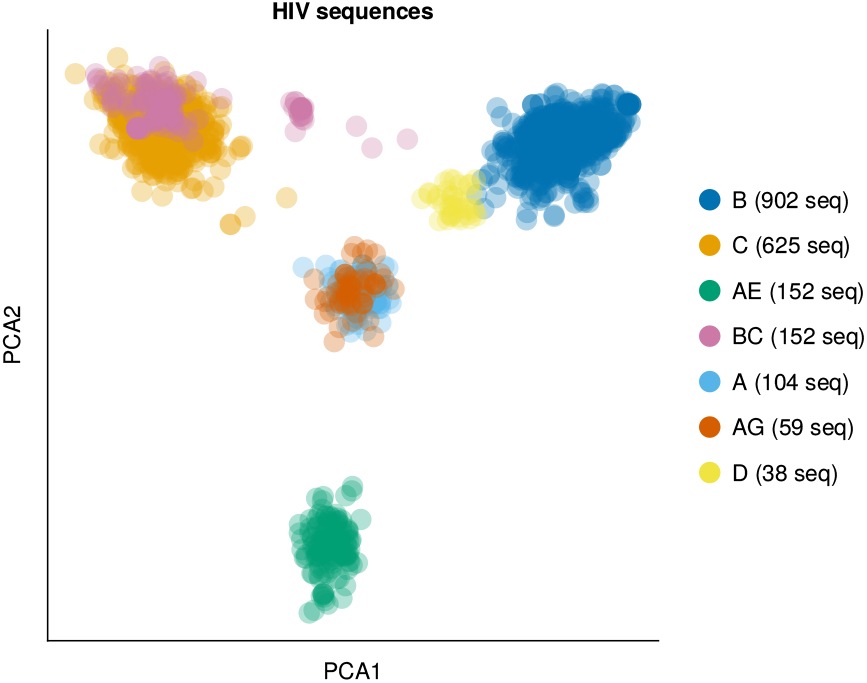
Distribution of HIV sequences across multiple subtypes in the first and second principal modes of the genetic covariance matrix. Each point in the cloud represents a viral sequence projected onto the principal modes of the covariance matrix derived from the one-hot encoded sequences. These principal modes yield projected sequences that are also used to learn the neutralization values. In this visualization, we present only subtypes that include more than 30 viral sequences that are used in our training data. Subtypes such as B, C, A, and AE are mapped to distinct regions in the space.

**Supplementary Fig. 3.**
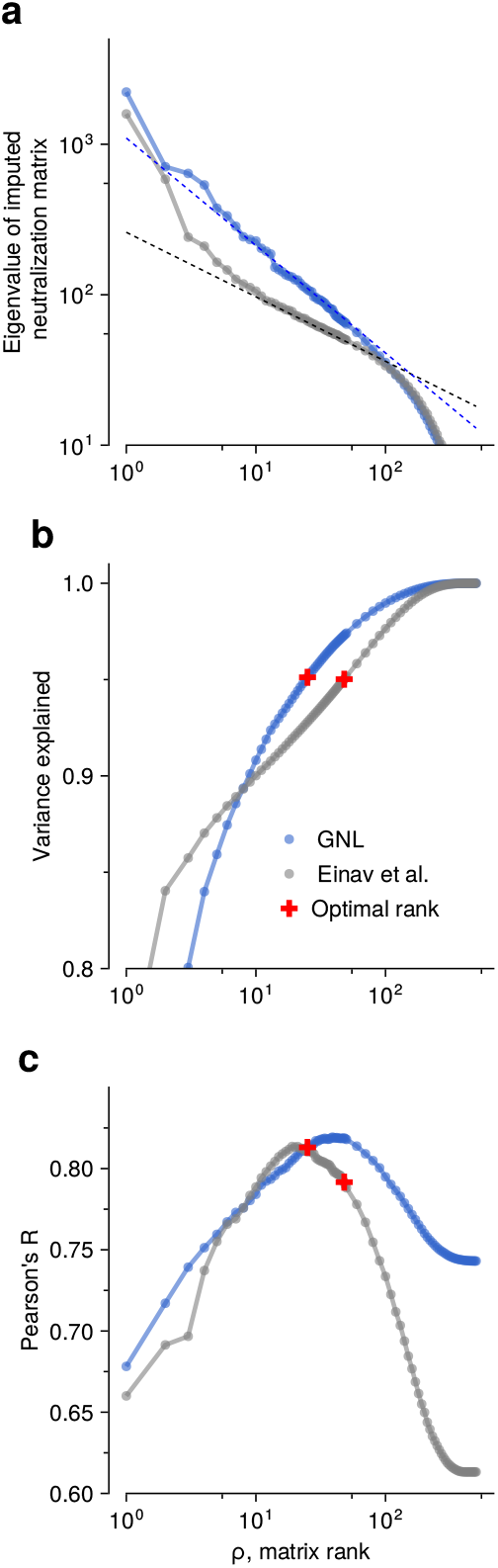
Variance explained and correlation between predicted and true neutralization values as a function of rank. As a typical example, we set the fraction of observed data as 80%, and the validation values were withheld uniformly at random from the CATNAP dataset. **a**, Eigenvalues for the filled neutralization matrices in the GNL and Einav et al. methods. The intermediate ranks of eigenvalues roughly follow a power law. Dashed lines show the power law fit with exponents of −3*/*7 and −5*/*7, respectively. **b**, Profiles of the variance explained for GNL and Einav et al. methods. The “optimal” matrix rank values, defined as the minimum rank at which the explained variance exceeds 95%, are 20 and 45 for the GNL and Einav et al. methods, respectively. **c**, Pearson’s R between true and predicted neutralization values shown as a function of rank. Including additional eigenmodes ultimately reduces predictive power. The rate of decrease in Pearson’s R using the GNL method is more gradual than that of Einav et al., and it maintains higher values.

**Supplementary Fig. 4.**
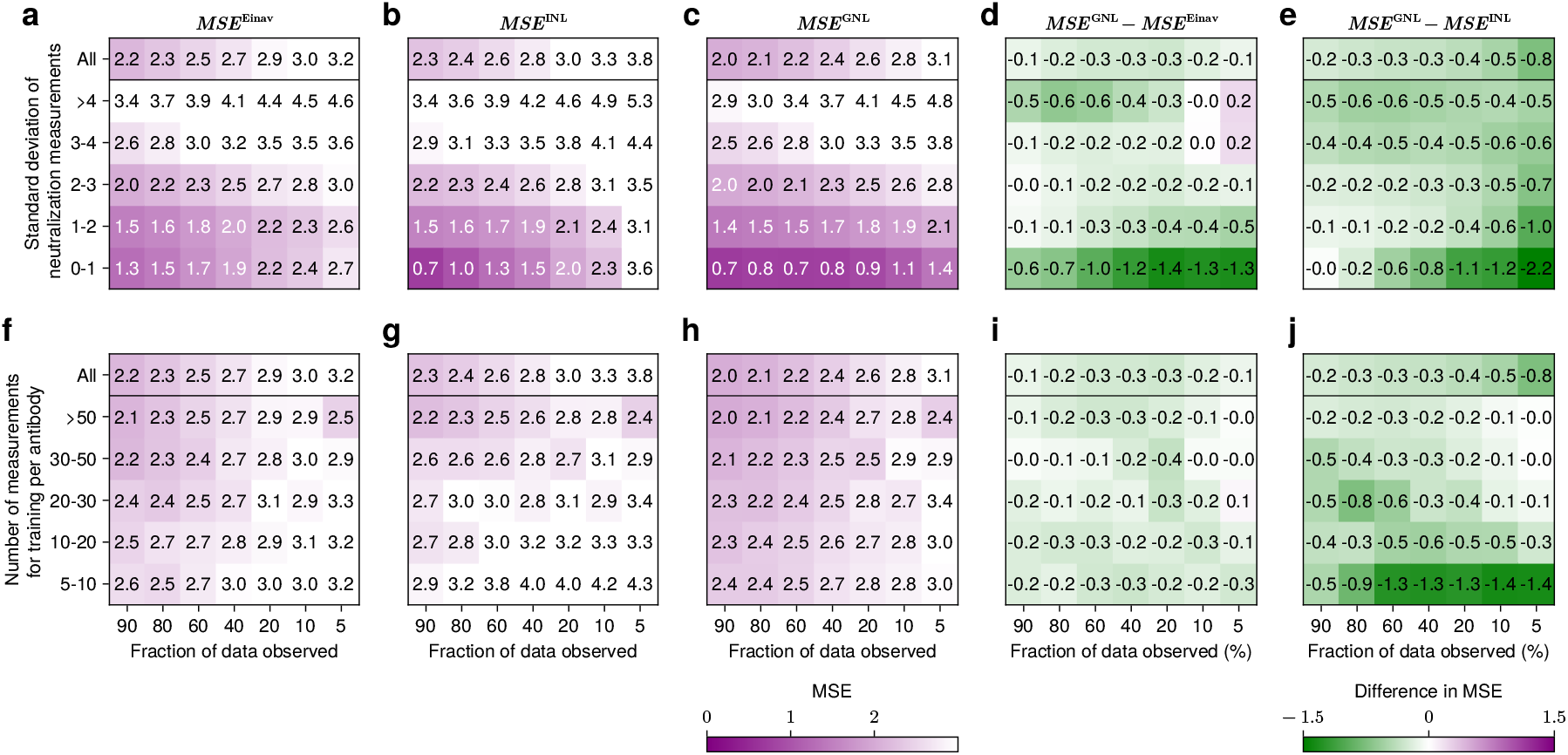
**a–b**, MSE values across seven data observation fractions (5% to 90%) and six classes of the standard deviation of neutralization measures, for the Einav et al. method, Independent Neutralization Learning (INL), and the GNL model. **d–e**, Differences in MSE values between GNL and the other methods. GNL outperforms INL for antibodies with higher variability in neutralization and for lower fractions of observed data (**e**), suggesting that antibody grouping mechanisms aid in learning from variable cases. **f–g**, MSE values across different numbers of training measurements and (**i–j**) their corresponding MSE differences. The grouping method benefits antibodies with fewer observed measurements, suggesting the advantage of grouping for sparsely observed antibody cases (**j**).

**Supplementary Fig. 5.**
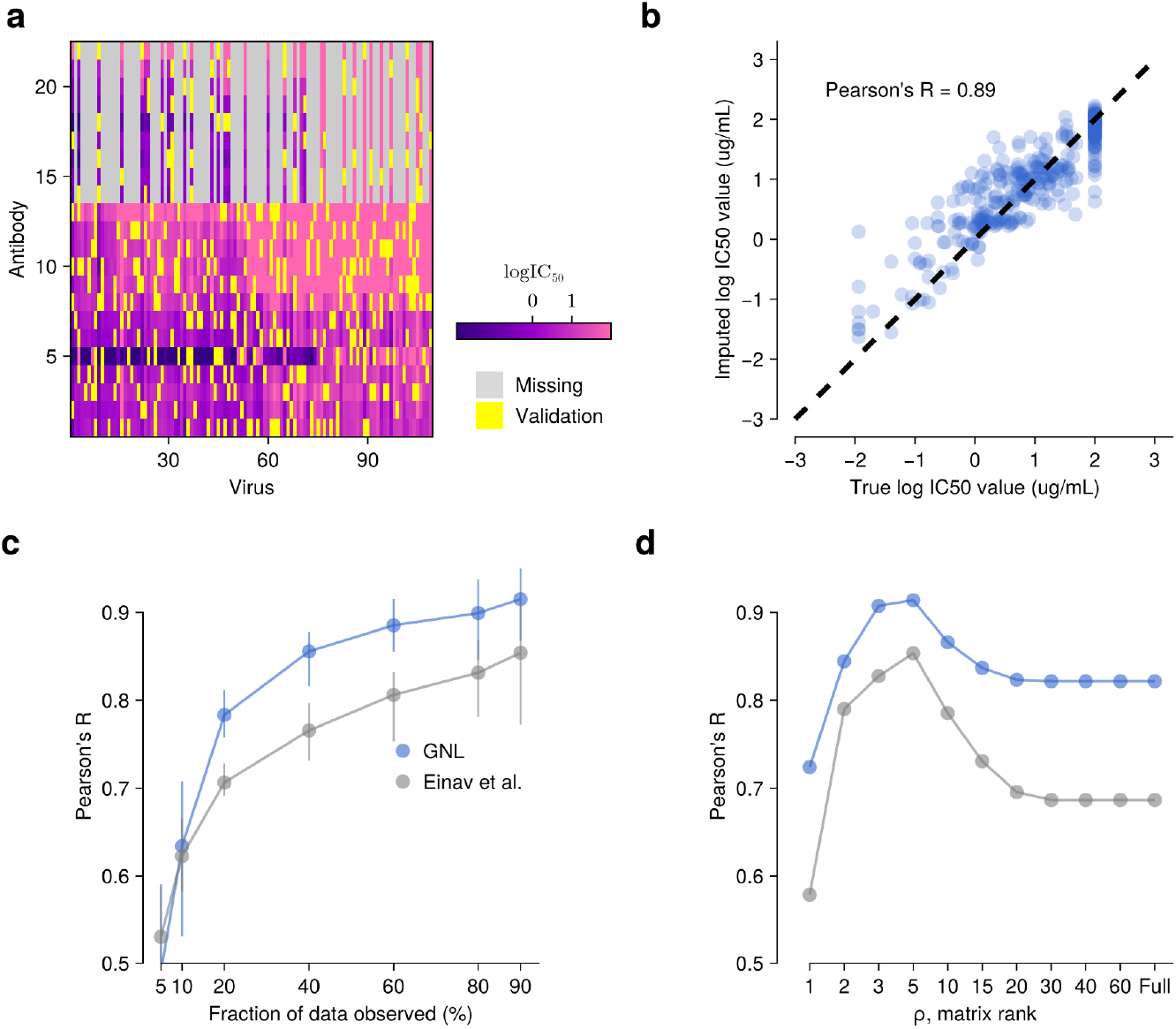
Overall neutralization imputation performance for the individual host. **a**, Neutralization values are represented with purple, gray, and yellow colors, corresponding to observed, missing, and withheld values for validation, respectively. The withheld elements are chosen uniformly at random across antibodies and viruses. In this analysis, 80% of the total available data is observed. **b**, A comparison of the true withheld neutralization values and the imputed values is shown on the x- and y-axes. Pearson’s R value is 0.89. **c**, Accuracy Dependency on the Fraction of Observed Data. The overall accuracy of the GNL method is higher than that of the Einav et al. method (provided by Einav et al. ^37^) as the fraction of observed data increases. **d**, Dependency of accuracy on matrix rank *ρ*. The R values of the GNL method are consistently higher than those of the Einav et al. method across most rank values.

**Table 1.** Model hyperparameters. Summary of key parameters, their meaning, suggested values, and guidelines for adjustment based on data characteristics.

| Notation | Meaning | Value or how to adjust it |
| --- | --- | --- |
| $\rho$ | Optimal neutralization matrix rank | Optimized as the minimum integer $\rho$ satisfying $\sum_{i=1}^{\rho} \sigma_i^2 / \sum_{i=1}^{\rho_{\max}} \sigma_i^2 > 0.95$, where $\sigma_i$ is the $i$-th singular value of the neutralization matrix. |
| $K$ | Maximum number of antibody groups | Use $K = 100$ for CATNAP. Adjusted to ensure well-distributed clusters while maintaining similarity within each antibody group. Results were insensitive in the range $K \in [50, 200]$. |
| $K$ | Max antibody groups (Intrahost) | Use $K = 5$ for intrahost neutralization data. |
| $d_{\max}$ | Max dimensions of projected sequences | Should be based on the maximum number of neutralization values across all antibodies. |
| $d$ | Dimensions of projected sequences | Optimized based on the number of viruses for which at least one antibody in each group has a neutralization value. |
| $d_k$ | Antibody group-specific dimension | Can be optimized depending on variation in neutralization profiles within each antibody group. |
| $\lambda$ | Regularization on regression parameters | We use $\lambda = 10$ for all conditions. Results were insensitive to values in the range $\lambda \in [5, 30]$. |
| $\lambda'$ | Similarity regularization between model parameters $\theta_\mu$ and $\theta_\nu$ | We use $\lambda' = 5$. This value was also insensitive in the range $\lambda' \in [1, 20]$. Note that $\lambda$ and $\lambda'$ should be jointly tuned. |
| $p$ | Exponent controlling similarity based on observation count | We use $p = 0.2$. This value was robust within the range $p \in [0.1, 0.4]$, but may need adjustment depending on the neutralization dataset. |

**Table 2.** Mean Pearson’s R values based on cross-validation. The mean cross-validation-based Pearson’s R (CV-R) values for each antibody type are listed in the table, with individual CV-R values shown in **Fig. 3** of the main text. The mean values were calculated only for bnAbs with sufficient data to allow SLAPNAP to run. For V1V2, V3, CD4bs, and MPER bnAbs, the mean CV-R values of GNL are often more than 10% higher than those of other methods. The overall mean CV-R, averaged across antibody types, indicates that GNL achieves the highest accuracy, with approximately a 10% improvement.

| Type of bnAbs | Einav et al.<sup>37</sup> | SLAPNAP<sup>40</sup> | GNL (our method) |
| --- | --- | --- | --- |
| V1V2 | 0.33 | 0.53 | **0.63** |
| V3 | 0.51 | 0.55 | **0.63** |
| CD4bs | 0.52 | 0.46 | **0.58** |
| Fusion peptide | 0.32 | **0.55** | 0.52 |
| Subunit interface | 0.21 | **0.43** | 0.34 |
| MPER | 0.65 | 0.59 | **0.75** |
| Total | 0.46 | 0.51 | **0.60** |

